# Genotypic and phenotypic diversity of the multidrug-resistant *Mycobacterium tuberculosis* strains from eastern India

**DOI:** 10.1101/2022.04.11.487831

**Authors:** Arup Ghosh, Himadri Bal, Viplov Kumar Biswas, Dasarathi Das, Sanghmitra Pati, Sunil Raghav

## Abstract

*Mycobacterium tuberculosis* (Mtb) poses a great challenge to human health and wellbeing and hinders economic growth of a region. India along with other south east Asian countries are known as high Tuberculosis burden countries. Adoption of whole genome sequencing in studying genetic diversity, evolution, transmission pattern and drug resistance development provided a great opportunity for developing and improving diagnostic and therapeutic approaches. In our study we have sequenced 118 Mtb whole genome from North East(NE) and Odisha as a representative of the diversity in eastern region of India for the first time. We observed high prevalence of multi-drug resistant(MDR) lineage-2(n=52) strains in NE whereas presence of mostly lineage-1(n=30) & 3 (n=11) strains in Odisha. The MDR strains from Sikkim demonstrated similar resistance profile of fluroquinolones and pair-wise SNP distances showed presence of local transmission clusters. We also detected significant enrichment of short INDELs in MDR samples in contrast to drug susceptible samples. This study provides molecular level insight into Mtb strains of eastern region in comparison with Indian and global perspective.

## Introduction

The COVID-19 pandemic has disrupted progress made for tuberculosis(TB) diagnosis in recent years and reduced access to TB diagnosis and treatment. The most visible impact observed in the global drop in the number of newly diagnosed and reported TB cases. A sharp decline of 18% observed from the data collected up to June 2021 (1). WHO Global Tuberculosis Report 2020 states that 26% of new TB cases are from India, which is the highest number among high TB burden countries followed by China with 9% of new cases reported. Although the number of new cases globally was lower than the 2017 report, there was only a marginal change observed in new cases from India(2). Also, India has the second-highest incidence of multidrug resistance (MDR) cases with the highest mortality rate (2). Although the drug-susceptible cases had a higher treatment success rate, the MDR and XDR TB cases had a treatment success rate of only 54% and 30%, respectively (3). The major challenge with the current TB diagnostic strategies are the time required for culture-based tests (3-6 weeks) and commercially available molecular diagnostics fail to account for novel compensatory mutations leading to drug resistance development (4, 5).

From the previous whole-genome sequencing reports and molecular dating (spoligotyping) studies it has been observed that the northern part of India is dominated by Lineage-3 whereas southern states show a prevalence of Lineage-1, central India has the presence of CAS & EAI, eastern states show a mixed diversity of lineage and the north-eastern states has a very dominant presence of Beijing lineage (6–9).

A genome-wide analysis study as a part of TB-ARC India project first time shed light on the lesser-known genetic diversity of the Mtb strains in the southern part of India (5). This study was conducted using 223 clinical isolates of which ∼15% strains were resistant for at least one drug when determined using phenotypic and genotypic drug susceptibility testing (DST) methods. In other studies, also where a lesser number of drug-resistant strains were investigated, the commercially available tests failed to detect the novel mutations present in Indian strains thereby resulting in a false negative outcome (6). A whole-genome study from The Foundation for Medical Research Mumbai also demonstrated the occurrence of novel mutations associated with drug resistance development(8). Recently another study with 200 *Mycobacterium tuberculosis* (Mtb) whole-genome sequences from ICMR-Jalma consisting of 91 MDR samples collected from north India identified novel resistance-associated mutations that are not used in any available molecular detection tests (7).

Whole-genome sequencing in recent times has provided us with a better understanding of drug resistance phenotypes and genomic diversity of Mtb globally (10, 11). Prediction of phenotype resistance utilizing genomic variants for some of the first-line drugs and widely used second-line drugs have shown promising results (12–14). In the case of India, the genomic surveillance of Mtb has been very limited and studies discussed earlier showed the presence of novel resistance-associated genotypes that can impact therapeutic outcomes (5, 7, 15). The current molecular tests used for rapid determination of resistance phenotypes mostly represent genotypes observed in global Mtb strains and are mostly dominated by European and American strains. In the case of India previous studies have shown that the resistance phenotypes vary at the regional level based on the lineage prevalence. It has been observed lineage 2 strains have a higher MDR rate in comparison to lineage 1 or 3 strains (11, 16, 17). Although whole genomic studies representing north and south India have been undertaken in recent years, such detailed genomic profiles are missing for eastern and north-eastern regions.

In our study, we have sequenced 118 culture-positive *M. tuberculosis* whole genomes which include 7 follow-up samples collected from Sikkim, Meghalaya, and Odisha from 111 patients. The samples collected from the northeast were composed of MDR, XDR samples as Fluoroquinolone resistance is highly prevalent in the region, whereas samples from Odisha mostly consist of drug-susceptible samples. Through this study, we expanded the understanding of the lineage diversity of the northeast and eastern region. We extensively studied the genomic diversity of Fluoroquinolone resistant Lineage 2 strains along with Lineage 1 and Lineage 3 strains of eastern India. We also examined the transmission patterns of MDR strains in the northeast and their mutation acquisition patterns using follow-up samples. Finally, we examined the performance of drug resistance phenotype prediction using a known set of mutations.

## Results

In this study, we carried out whole-genome sequencing of 118 tuberculosis isolates cultured using mycobacteria growth indicator tube (MGIT) from North East (n=70, including follow-ups) and Odisha(n=48) representing north-east and east India collected between February 2017 to June 2020. Out of the 118 samples, there were 7 follow-up isolates belonging to five patients collected between April 2018 and September 2019 from Sikkim and were removed from the comparative analysis. Out of the remaining 111 samples, 9 samples didn’t pass the MTBC percentage filtration(>=80%) and pruned from further analysis.

The isolates collected from Sikkim and Meghalaya were all drug-resistant samples sent to the RMRC-Bhubaneswar reference laboratory after the Cartridge Based Nucleic Acid Amplification Test (CBNAAT) test showed presence of Rifampicin resistance associated variant. Majority 51/54 of the isolates collected from the northeast were multi-drug resistant (MDR) samples and 3/54 were classified as extensively drug-resistant (XDR). The samples from Odisha were 39/48 drug-susceptible and except 9/48 samples out of which 2/48 classified as mono resistant and 7/48 as MDR.

The median age of patients enrolled in this study is 32.0±17.01 years consisting of 57 Males, 44 Females sex, for 1 individual gender was not specified (Table -1, Supplementary Figure 1A). Information related to HIV status, smoking or alcohol consumption were not reported. We further curated two major whole-genome datasets (n=423) published from India with respective DST information (7, 18). For the South Indian samples, we collected the variants call files and phenotypes from the PATRIC database, and North India cohort samples were reanalysed following our sequence analysis criteria 471 (Sikkim-odisha n=102, curated n=369) sequences from India were used for comparative analysis(19).

**Table 1:** Demographics of patients enrolled for the study, drug resistance phenotype of sequenced samples (excluding follow-up samples) and lineage distribution (detected by variant calls against published panels)

| *Age (yr)* | *Count* | *Percentage (%)* |
| --- | --- | --- |
| <30 | 41 | 40.20 |
| >30 | 61 | 59.80 |
| **Sex** |  |  |
| *M* | 57 | 55.88 |
| *F* | 44 | 43.14 |
| *Undisclosed* | 1 | 0.98 |
| **Region** |  |  |
| *North-East* | 54 | 52.94 |
| *East* | 48 | 47.06 |
| **Resistance (WGS)** |  |  |
| *DS* | 39 | 38.24 |
| *MONO* | 2 | 1.96 |
| *MDR* | 58 | 56.86 |
| *XDR* | 3 | 2.94 |
| **Lineage** |  |  |
| *Lineage-1* | 31 | 30.39 |
| *Lineage-2* | 56 | 54.90 |
| *Lineage-3* | 12 | 11.76 |
| *Lineage-4* | 3 | 2.94 |

**Table 2:**
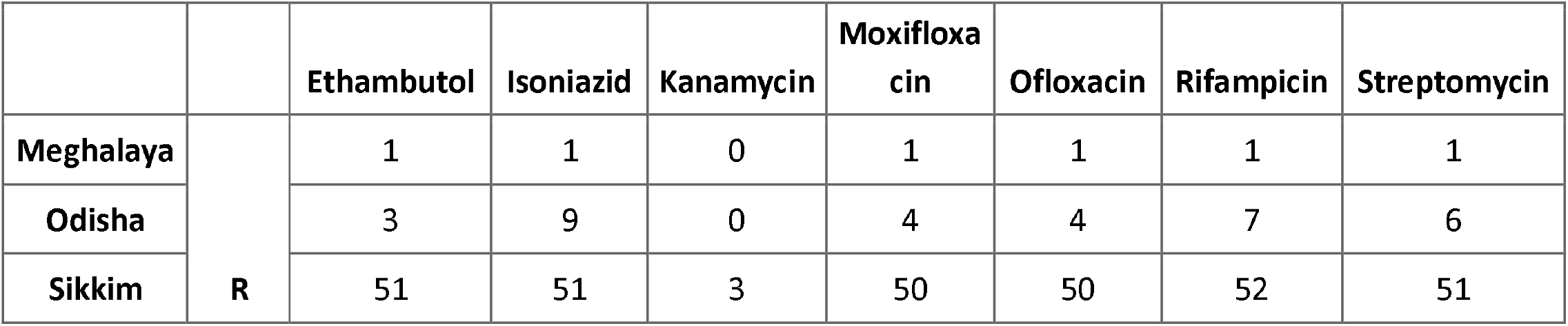
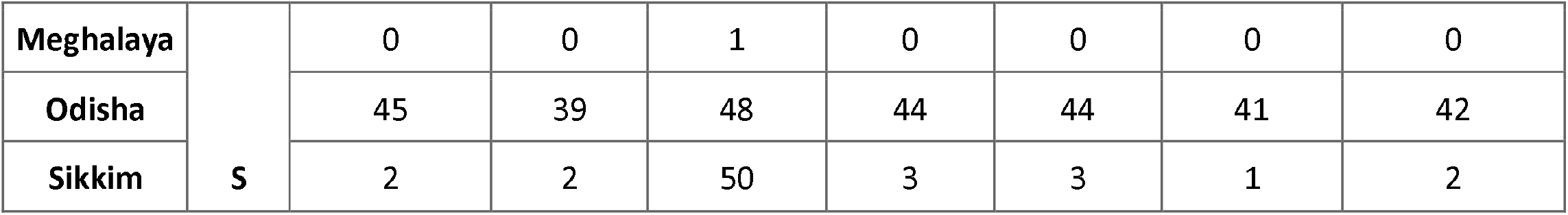
Summary of drug resistance phenotypes for seven drugs predicted by Mikrobe predictor tool(26), grouped by place of sample collection. R = Drug resistance & S = Drug susceptible

|  |  | Ethambutol | Isoniazid | Kanamycin | Moxifloxacin | Ofloxacin | Rifampicin | Streptomycin |
| --- | --- | --- | --- | --- | --- | --- | --- | --- |
| Meghalaya | R | 1 | 1 | 0 | 1 | 1 | 1 | 1 |
| Odisha |  | 3 | 9 | 0 | 4 | 4 | 7 | 6 |
| Sikkim |  | 51 | 51 | 3 | 50 | 50 | 52 | 51 |
| Meghalaya | S | 0 | 0 | 1 | 0 | 0 | 0 | 0 |
| Odisha |  | 45 | 39 | 48 | 44 | 44 | 41 | 42 |
| Sikkim |  | 2 | 2 | 50 | 3 | 3 | 1 | 2 |

Using the QuantTB algorithm for mixed infection detection using whole-genome sequences we observed similarity of NE and Odisha strains with other Indian or South East Asian strains belonging to same lineages and only one strain detected per sample (20).

### Genome-wide variations shed light on *Mycobacterium tuberculosis* genetic diversity in North-East and Odisha

We detected a total of 12926 high-quality single nucleotide variants, 825 short indels with lengths ranging from 29bp (max insertion) to 48bp (max deletion length) (Supplementary plot 1C) and a total of 2878 large deletions (Supplementary plot 1D). Effect of variants in respective genes are predicted using snpEff tool (Figure 1D). Using the high-quality single nucleotide variants we determined lineages for all the *Mtb* samples with Coll 2014 SNP barcodes and fast lineage caller tool (21, 22). We observed a drastic difference between the lineage distribution of Mtb in the northeast (NE) region and Odisha. Lineage-2 also known as Beijing lineage has a very dominant presence in NE 62/64 whereas Lineage-1 also known as Indo Oceanic lineage is more prominent 30/48 in Odisha (Figure 1B). The prevalence of Beijing lineage in NE regions is coherent with a recent spoligotyping-based survey conducted in Sikkim, the authors observed 62.41% occurrence of *Beijing* strains in the region (23). Although lineage-1 is very dominant in Odisha the region also sees a prevalence of lineage-3 11/48 also known as Central-Asian lineage followed by Beijing and lineage 4 (European/American) strain (Figure 1B). To further understand the most affected areas we checked the distribution of lineages among different districts (Supplementary table1) of these two states and found that most of the samples are from eastern region 51/64 in case of Sikkim. The southern districts of Odisha are mostly affected by lineage-1 whereas central and eastern districts show a mixed presence of all three lineages. As there were no previous large scale genomic or spoligotyping surveys were not available from Odisha we assume that the genetic diversity of Mtb in this region is heavily influenced by neighbouring regions a high prevalence of lineage 1,2 and 4 (6, 7).

**Figure 1.**
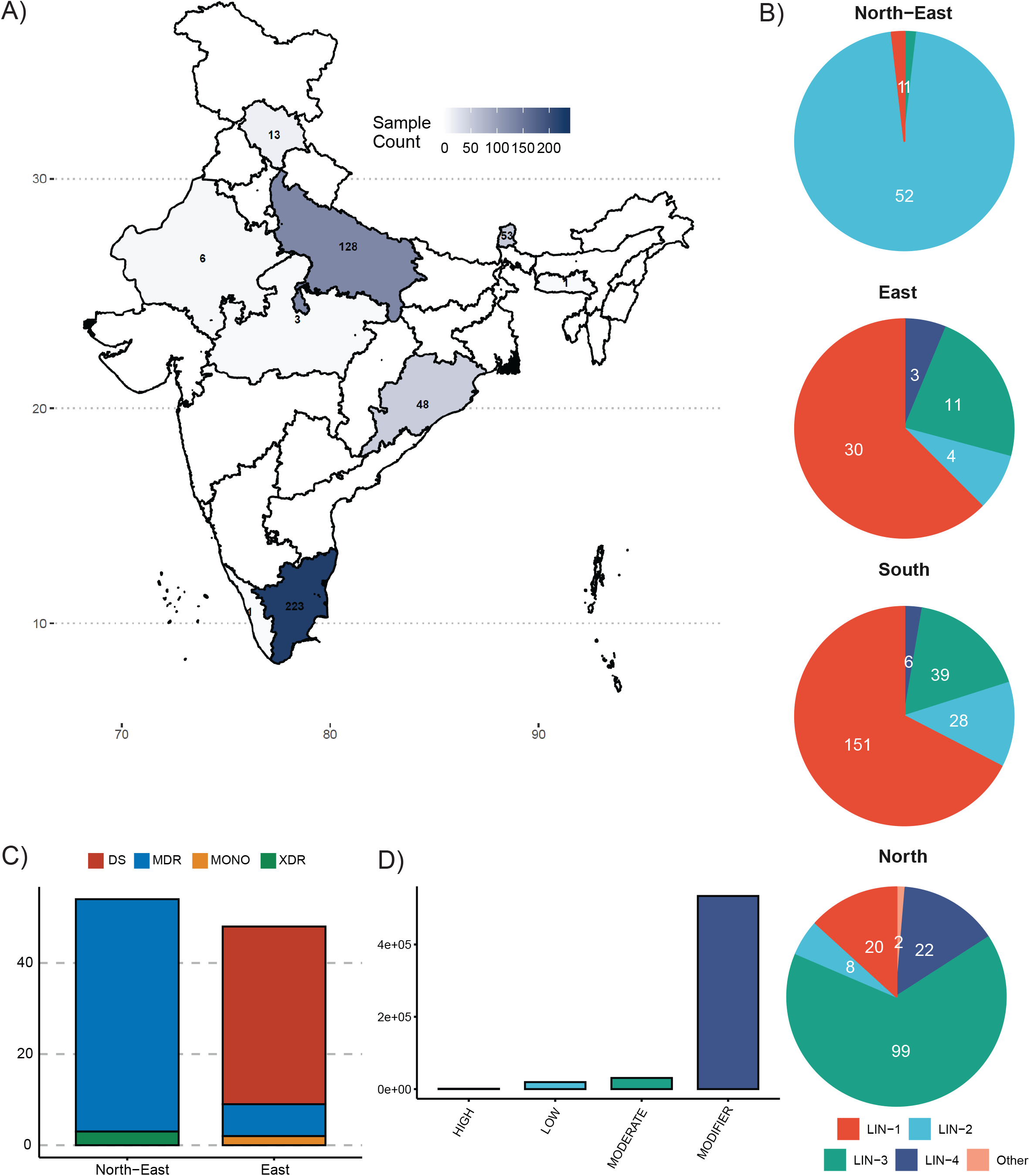
A) M. tuberculosis whole genome sequence samples from our study and collected from other studies for comparative analysis. B) Distribution of Mtb lineages across different regions with number of samples. C) Drug resistance phenotype of samples from our study represented by stacked bars and groups by respective regions. D) Effect of variants on corresponding gene predicted using snpEFF.

To understand the genetic diversity in context with other strains prevalent in India we constructed phylogenetic tree of all 102 samples from our study along with two previously published studies depicting the lineage diversity in south and north India.

### Phylogenetic classification and genetic similarity with other regional and global strains

Single nucleotide variants from 471 samples consisting of previously published and newly sequenced *Mtb* whole genomes from India were used to create neighbor-joining phylogenetic tree with 1000 bootstraps(24). The tree represents the genetic diversity of Mtb across India represented by their geographical location and drug resistance phenotypes (Figure 2). The drug resistance *Mtb* samples from Sikkim were divided into two sub-clusters while coinciding with other lineage 2 samples from Tamil Nadu and Uttar Pradesh. One of the Sikkim sub-clusters branches show very small branch lengths from the nearest node, indicating presence of local transmission from a most recent common ancestor (MRCA) of the strain (Figure 2).

**Figure 2.**
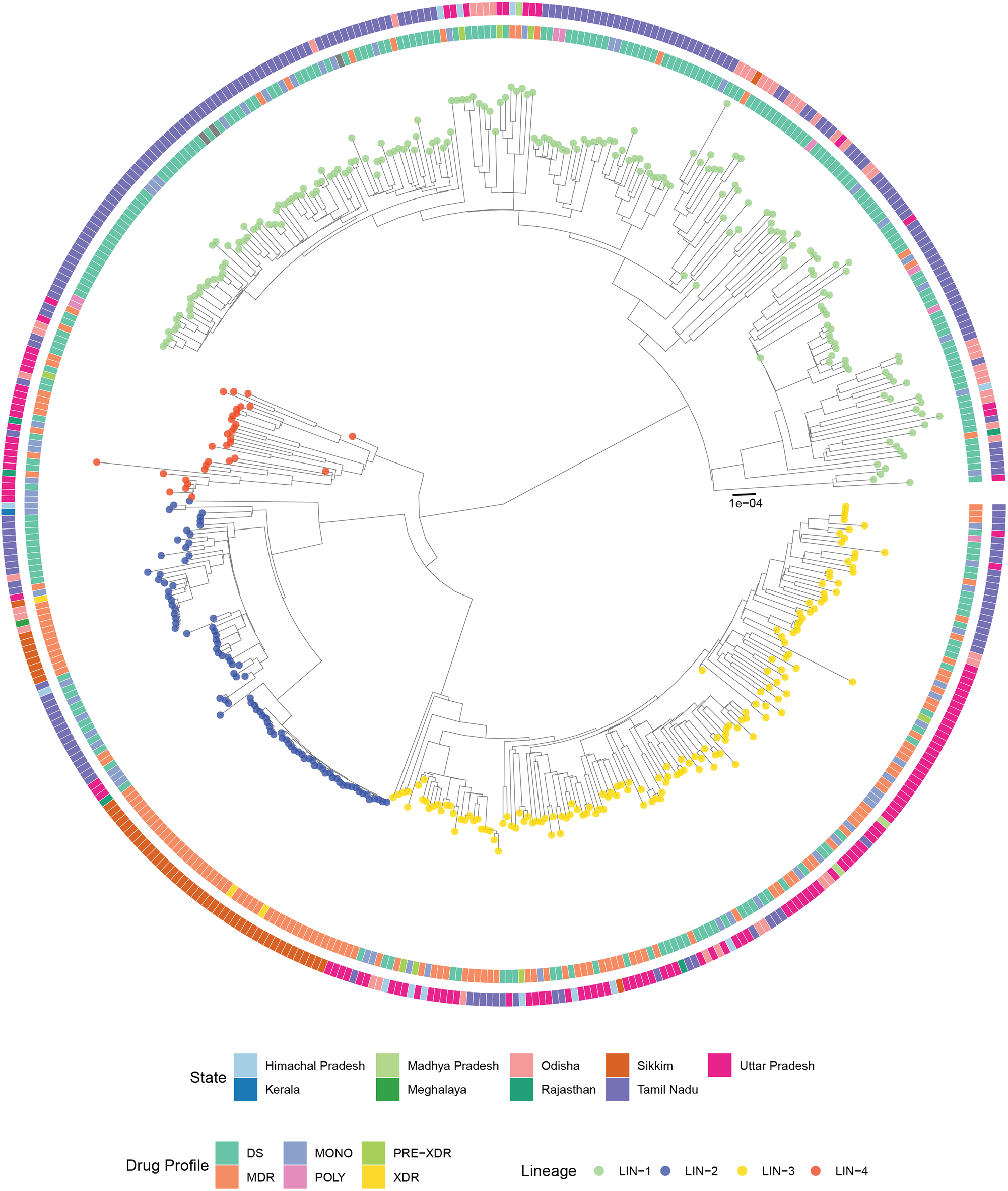
Neighbor joining tree representing M. tuberculosis isolates from North, South, North-East and East India isolates, with 1000 bootstraps (Outgroup M. canetti not shown). The node colour represents isolate lineage, inner circle shows corresponding drug resistance phenotype and outer circle coloured based on the isolates geographical location (divisions/states).

The samples from Odisha were divided between lineage 1 and lineage 3 clusters and supporting our previous hypothesis they grouped with representative strains from North and South India respectively (Figure 2). The lineage 4 samples from Odisha clustered with similar strains from Uttar Pradesh and Rajasthan (Figure 2A).

To further extend the understanding of genetic diversity of the strains in global context we performed principle component analysis(PCA) of the 471 samples along with a set of 2500 global *Mtb* samples curated from GMTV dataset (25). Using principal component 1 and 2 ∼87% of variance was explained (Figure 3A). The strains from India grouped among the global lineages based on their respective lineages but showed some deviation in case of lineage 2 samples from Sikkim and lineage 3 samples from North India (Figure 3A).

**Figure 3.**
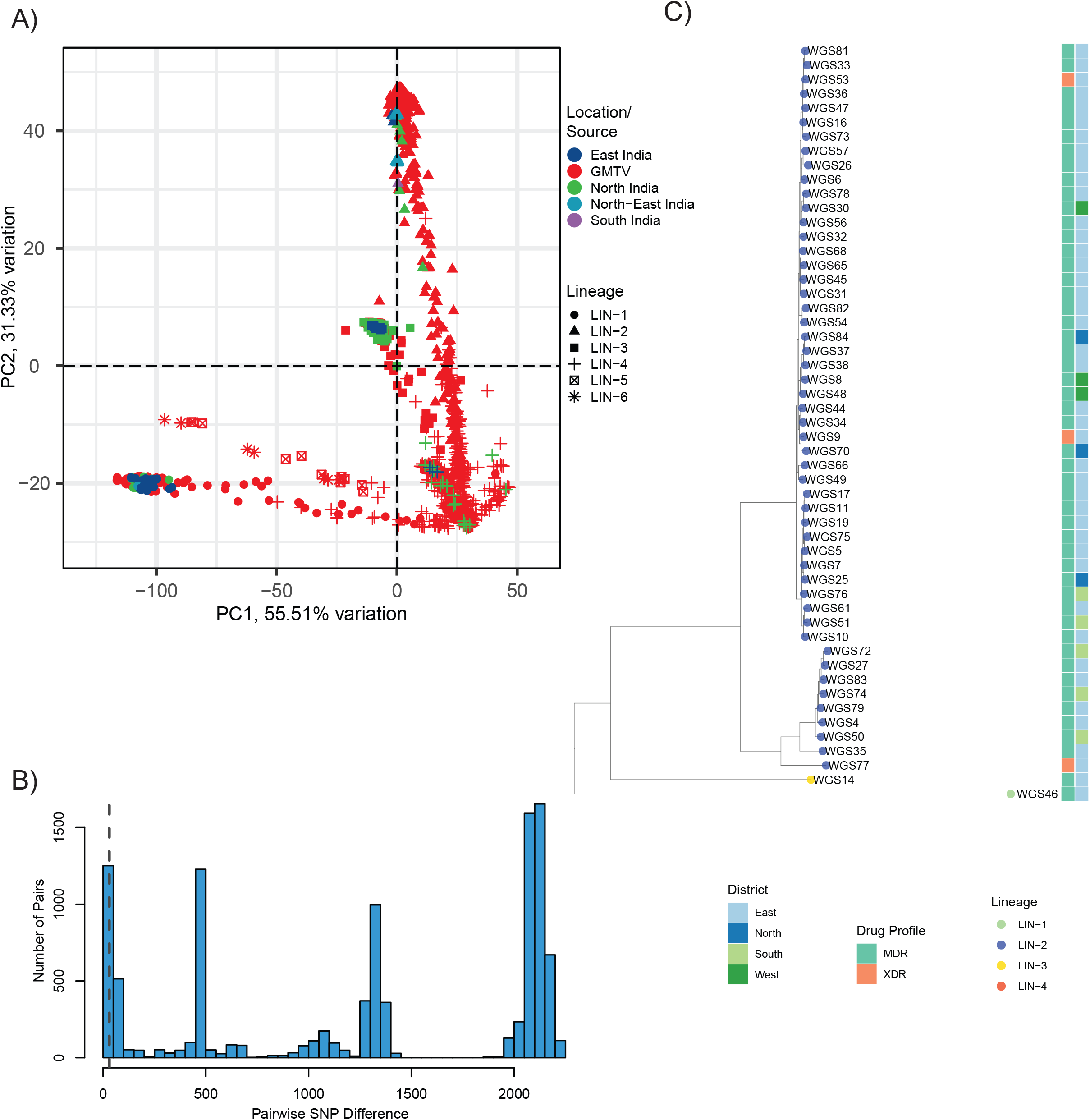
A) Principle component analysis of Indian tuberculosis strains with GMTV strains, shape of the dots are used to represent the lineage and colour represents the dataset/strains source. B) Distribution of pair-wise SNP distances of North East and Odisha sequences, dashed line represents 35 SNP differences among the pairs. C) Phylogenetic tree representing the Sikkim sample local transmission clusters with their drug resistance phenotypes, lineage and collection district.

The two clusters of drug-resistant *Mtb* samples from Sikkim observed in the phylogenetic clustering of the samples, indicated the presence of local transmission clusters in the region and to further investigate we utilized SNP distance based methods.

### Local transmission of drug resistance Beijing strain in Sikkim

In accordance with our findings from phylogenetic tree we calculated pair-wise distance among all the sequenced samples from our study (n=102) and plotted the distribution of pairwise SNP distances. We observed two group of lineage 2 sequences (n=42,9) from Sikkim showing pairwise SNP distance of less than 50 in contrast to other strains showing at least 100 or more dissimilar SNPs among them (Figure 2B). We clustered the pairwise SNP information using a hierarchical clustering method and observed to distinct clonal clusters of drug-resistant samples collected from Sikkim (Supplementary Figure 2). To further resolve the genetic similarity among the strains we created a phylogenetic tree of all 53 samples from the region and included their district level information. The resultant phylogenetic tree represented the main cluster (n=42) consisting strains mostly from east Sikkim with 3/42 strains each from north and west Sikkim. The smaller cluster (n=9) also represented sequences from east district along with 3/9 strains from west district (Figure 3C).

### Drug resistance phenotype of North East Indian strains matches with global Beijing strains with high Fluoroquinolone resistance burden

One of the main goals of our study was to understand the genomic variations leading to high fluoroquinolone (namely moxifloxacin(both low and high concentrations), levofloxacin, and ofloxacin) in the state of Sikkim. The abundance of multi-drug resistance strains among Lineage 2(Beijing) lineage is very high. Out of 53 drug-resistant samples collected from Sikkim 51 (96.22%) belonged to Lineage 2 (sublineage 2.2.1) of which 48 (94.11%) belonged to the multi-drug resistance category and 3 (5.88%) in the extensively drug-resistant category. As per the scope of the national tuberculosis surveillance program, only a limited number of drugs were tested for DST but with the help of whole-genome sequencing data, we predicted the phenotypes for 11 drugs used for the treatment of tuberculosis. We observed a high prevalence of rifampicin resistance followed by isoniazid, ethambutol, streptomycin, and fluoroquinolone group of drugs in Sikkim. On the other hand in Odisha isoniazid resistance is more prominent in comparison to other groups of drugs.

We compared the predicted results with our first-line and second-line LPA assay data and two culture-based DST results. In the case of rifampicin the prediction was able to explain 101/102 cases, but in the case of kanamycin and capreomycin resistance there were 6/99 unexplained cases. In the fluoroquinolone category the results overlapped in 95/98 cases, thus mostly explained by known genomic changes. To get a better perspective of variant accumulation loci we ranked the genes based on the number of non-synonymous mutations present in the gene body followed by functional segregation. The top biological processes came up as metabolism and respiration, information pathway, lipid metabolism, cell wall-related process, and virulence (Figure 4A). All of these pathways are known to be directly associated with drug resistance phenotypes and the changes in virulence-related genes might have a role in the increased pathogenicity of Beijing lineage, which is overrepresented in our dataset (Figure 4A)(27). When we looked into genes with highest number of non-synonymous variations acquired, genes from RNA polymerase and DNA gyrase coding family ranked in the top position (Figure 1B). Followed by our observation we checked for the presence of known drug resistance associated loci and their frequencies (count), the results were coherent with prediction outcomes (Figure 4C). Although the functional impact if INDELS has been less studied in the context of drug resistance association and other functional implication, number of INDELS detected in MDR samples were significantly higher in contrast with drug susceptible sample and the XDR samples also show an increased pattern (Figure 4D). The MONO resistant sequences only had slight increase in the INDEL number in comparison with susceptible sequences.

**Figure 4.**
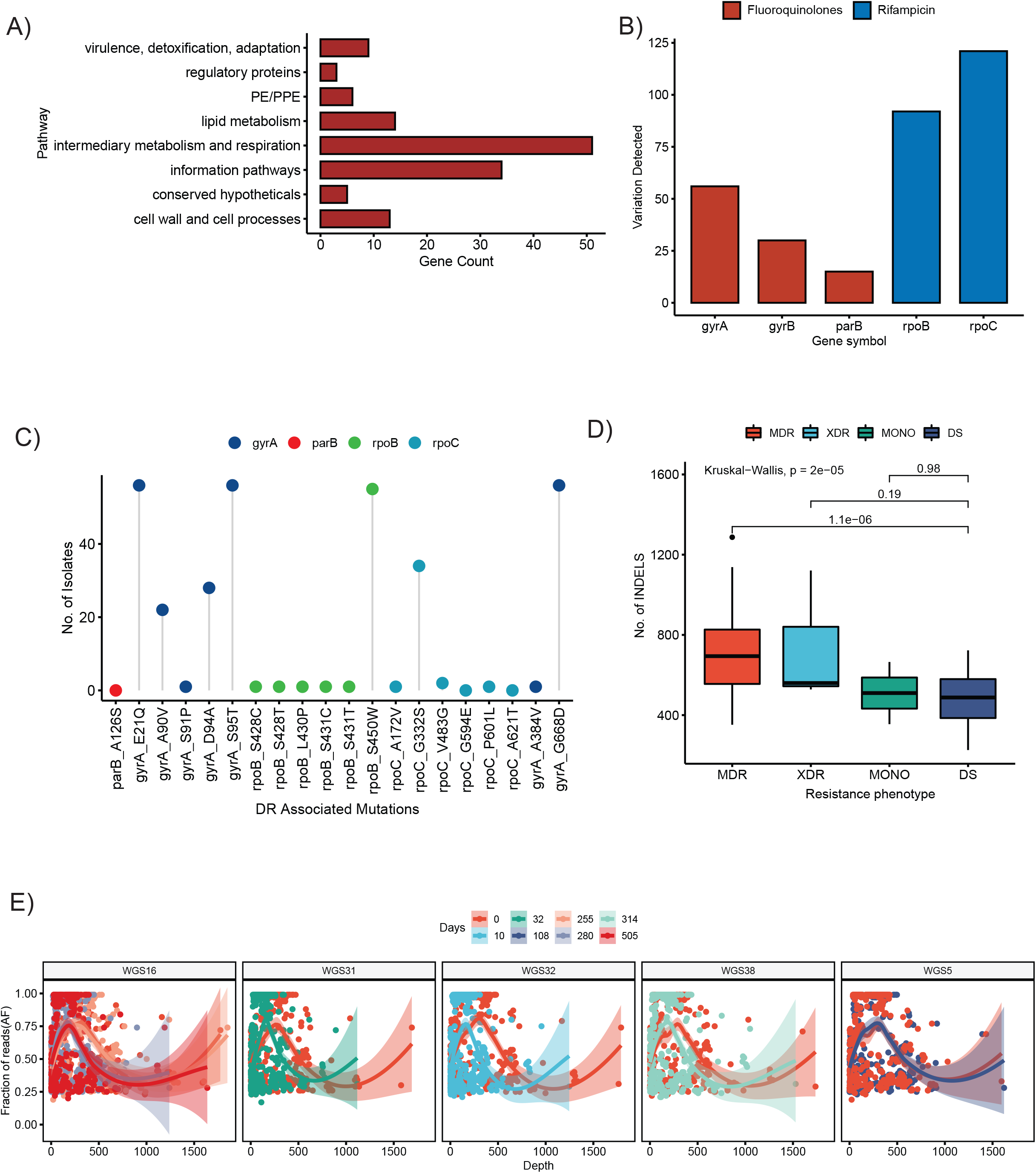
A) Functional categories of highly mutated genes (ranked based on number of non-synonymous mutations detected). B-C) Drug resistance associated genes and amino acid changes with respective mutation density and frequency in sequenced samples. D) Number of INDELs detected per phenotypic categories compared with susceptible samples compared using non-parametric test. E) Allele fractions of follow up isolates plotted against respective depth of coverage coloured by days post initial collection.

### In-host pathogen dynamics in contrast to resistance-associated loci

During the process of samples collection we also followed up with six patients from Sikkim. We collected a total of 7 isolates from five patients in an interval of 10 days to 1.5 years to examine if there is presence of in-host variations. Genetic variations fixed during the while the treatment regimen is ongoing sheds light on clonal selection of drug resistance Mtb strains and recently has been shown to have impact on host innate immunity and metabolism(28). The pairwise SNP distance among the same follow-up patients ranged between 69-105 (Median: 89.50) indicating that despite slow mutation rates (0.3-0.6 substitutions/per year) and lack of recombination some (22).To elucidate the presence of in-host variation overtime we grouped the follow up sequences with the primary strain and plotted the distribution of allele fractions (fraction of reads supporting an allele) against depth of coverage of the region (Figure 4E). Trendline in the data showed increased fractional representation of some of the variant positions although when we compared the allele fractions of known resistance associated loci, they were fixed (AF = 1) throughout the timeline (Figure 4E).

## Discussion

The focus of our study was to explore the genetic diversity of M. tuberculosis complex in North East and Odisha where no prior whole-genome sequencing studies have been undertaken. For the first time, our study shares insight about Mtb lineages in the state of Odisha and their genetic diversity. In addition, our study provides insight into the genetic basis of the high prevalence of multi-drug resistance Beijing strains, local transmission of fluoroquinolones resistant strains, and acquisition of resistance-associated mutations in North-Eastern states represented by Sikkim.

We observed a high occurrence of a localized patient-to-patient transmission pattern of MDR TB Beijing strains (<35 SNPs) in North East India. The contact-based transmission of MDR strains might be one of the reasons for the high abundance of fluoroquinolone resistance along with rifampicin resistant cases in the region. The cluster in East Sikkim indicates the probable presence of nosocomial transmission of Tuberculosis in the region, which might pose challenges to the ongoing mitigation programs and required to be taken into account for prevention programs (17). But Due to the lack of sufficient metadata and number of samples we were not able to determine whether the transmissions events took place in a healthcare centre or region-specific manner.

The North East region is known to have a very high burden of drug-resistant tuberculosis cases and a recent study accessing molecular diversity of MTBC in the regions showed a high prevalence of Beijing strains (∼62%) and higher occurrence of MDR phenotypes in the strain(29.9%) in comparison to other MTBC lineages(4.7%) (12, 18). Our observation is consistent with the previous findings additionally we observed an increased burden of fluoroquinolone resistance in the region as well (12). Although our isolates from Sikkim were enriched for rifampicin (CBNAAT) resistance phenotype the concurrent findings suggest our results are representative of NE region the dominance of Beijing lineage and multi-drug resistance (12).

Follow up samples collected from Sikkim in a period of ∼1.5 years depicted change in allele frequency of strains accumulated over the period of sample collection. With the limited number of samples we were not able to capture any resistance associated changes. But such studies can provide insight in to the initial acquisition of fluroquinolone resistance burden in the region as amplification of fluroquinolones and second-line injectables occurred in a very recent time(29).

On the other hand in Odisha we observe a very diverse representation of Mtb strains, all three lineages prevalent in India observed in this region. Lineage 1 is the dominant followed by lineage 3 and 2. The distribution of lineage has a similar pattern as the study conducted in Tamil Nadu, the cause might be the frequent migration of people between regions in search of livelihood. Although Odisha has less burden of drug resistance tuberculosis the number of reported cases are also less due to lack of health facilities in close vicinity, and poor lifestyle choices (30). Odisha is a tribal population dominated region and the community is scattered across dense forest areas or mountains making it very difficult to estimate the actual burden of tuberculosis. Studies performed in nearby state of Madhya Pradesh shared insight into high tuberculosis burden Saharia tribe(20.4%) a geographically isolated population of agriculture labours(31).

Detection of drug-resistant phenotype in a very early stage of diagnosis helps in the determination of treatment regimen, which is a major contributor in determining the treatment outcome. Due to the huge burden of new cases, sometimes confirmatory tests for second-line drugs resistance take up to 20-30 days after initial detection of Rifampicin resistance using rapid PCR-based techniques. Recent studies have demonstrated the reduced efficacy of GeneXpert MTB/RIF, Hain MTBDRplus, and Hain MTBDRsl in the detection of drug resistance phenotypes due to region-specific mutations that are not incorporated in the tests (19).

In the newly sequenced samples we were able to explain most of the drug resistance ceases with genotype-based predictions but for a small fraction of sequences, the prediction-based approach failed to account for outcomes. Majority of the phenotypic resistance has been explained by mutations in gyrase protein coding genes and rna polymerase subunit coding genes. A large proportion of Rifampicin and Fluroquinolone resistant(MDR) samples harboured gyrrA E21Q, D94A, G668D and A90V along with rpoB S450W, rpoC G332S amnino acid substitutions. For the cases where Mykrobe predictor failed to associate phenotype based on genotype information, depicts a lack of region specific resistance conferring features (26).

In summary large scale genomic studies are required to understand the dynamics of highly diverse Mtb population India to device better region specific diagnostics methods. Also large scale genome sequencing undertakings will help improving already available low cost molecular diagnostic tests. Using large scale association studies targeted panels can be developed for prediction of drug resistance phenotypes directly from sputum samples in short period of time using massively parallel sequencing approach.

## Methodology

### Sample collection and MGIT culture

The samples sequenced/used in this study were initially processed and cultured in National Reference Laboratory, Regional Medical Research Centre, Odisha. The study was conducted in accordance with recommended guidelines and safety procedures of ICMR-Regional Medical Research Centre, Odisha. All the subjects gave written informed consent in accordance with the Declaration of Helsinki.

The samples used in this study were randomly selected from a pool of CBNAAT tested known Rifampicin resistance samples from North East India and sputum samples collected from Odisha submitted for testing of the presence of *M. tuberculosis*. All the sputum samples were initially treated with N-acetyl-l-cysteine-sodium hydroxide (NALC-NaOH) for decontamination and inoculated on Lowenstein-Jensen (LJ) slants for primary culture and mycobacteria growth indicator tube (MGIT) subsequently. Once the MGIT tubes showed detectable growth levels of the bacteria they were kept in an incubator for another 2-3 weeks to achieve sufficient numbers of bacteria for DNA isolation.

### Drug susceptibility testing

The sputum samples were tested for drug resistance using four different WHO-recommended testing methods. The primary test was CBNAAT for checking the presence of Rifampicin resistance in primary collection centers, the results were further validated using drug resistance first line LPA testing for Rifampicin and Isoniazid (Genotype MTBDRplus version 2.0 Hain Lifescience GmbH), samples showing resistance were further tested for second-line LPA consisting of Fluoroquinolone resistance second-line injectables and Kanamycin (low) mutations (GenoType MTBDRsl by Hain Lifescience GmbH). Samples showing Fluoroquinolone and MDR phenotype were further tested for second-line drug susceptibility testing using 1% proportion DST method by liquid BD BACTEC MGIT automated culture challenged with Kanamycin (2.5 μg/ml), Amikacin (1.0 μg/ml), Capreomycin (2.5 μg/ml), Ofloxacin (2.0 μg/ml), Levofloxacin (1.5 μg/ml), Moxifloxacin (0.5 and 2.0 μg/ml) in a BD BACTEC™ system to check the growth of bacteria sufficient for conferring resistance. Samples showing Rifampin resistance were directly tested for second-line resistance using LPA as per NTEP diagnostic recommendations.

### DNA isolation and library preparation

The MGIT cultured samples were kept for another 2-3 weeks to get sufficient growth for DNA isolation. DNA from all the samples were isolated using a bead beating method using HiPurA™ (catalogue number: MB545) from HIMEDIA labs inside the Biosafety level 3 (BSL-3) facility of ICMR-RMRC, Bhubaneswar. The DNA quality of individual tuberculosis samples were checked using NanoDrop™ spectrophotometers and samples having a 260/280 ratio between 1.6-2.0 were selected for preparation of whole genome sequencing libraries. All sequencing libraries were prepared using Nextera XT DNA library preparation kit from Illumina, Inc with 1ng of input DNA in 0.2 ng/ul concentration. The prepared whole-genome sequencing libraries were pooled and sequenced using NextSeq 550 system with a 2 × 150bp paired-end layout.

### Quality control and classification of MTBC percent

To check the percent of Mycobacterium tuberculosis complex in each clinical isolates we ran Kraken2 (PMID) against a database containing archaea, bacteria, viral, plasmid, vector in RefSeq and human reference assembly (minikraken_8GB_20200312). The proportion of reads falling under MTBC were summarised using a recently published method (32). Samples having less than 80% of reads classified ad MTBC were removed from further analysis.

### Alignment and variant calling

The samples passing MTBC filtration criteria were mapped to the H37Rv (NC_000962.3) reference genome obtained from NCBI using BWA-MEM(33). Duplicate reads from all the samples were removed using Picard before calling variants. Single nucleotide variants and short insertion and deletions were called using Pilon with --fix all, breaks option. From pilon output sites with “PASS” filter were extracted and divided in SNV and INDEL calls (34). We further filtered all variants with minimum map quality (MQ) score of 30, minimum base quality (BQ) score of 20, minimum read depth of 10 or more than 80% reads supporting alternate genotype, and no insertion or deletions called in the region (IC=0 and DC =60). The filtered variants were then merged using bcftools merge and annotated using SnpEff(Supplementary plot 1D) with H37rv reference database(35, 36). For further analysis genotype matrix and annotations were extracted using bcftools and SnpSift(37).

Unfiltered variant call files for South India samples were directly downloaded from Broad Institute portal (https://olive.broadinstitute.org/collections/tb_india.1) and the variant positions were updated to NCBI H37Rv reference assembly using Picard LiftOver utility and CP003248.2 to NC_000962.3 conversion chain file generated by UCSC toolkit.

### Lineage calling

Lineage determination of all the samples were performed using Fast-line-caller from Farhat-lab(22). The tool takes vcf files as input and assigns lineages based on presence of pre-defined lineage determining mutations reported by multiple sources. We also predicted digital spoligotyping using SpoTyping tool and the predicted types were annotated by querying SITVIT database, both of the methods provided similar lineage calls and lineage nomenclature provided in Fast lineage caller with Colls set were considered in final analysis (21, 38).

### Phylogeny reconstruction

Phylogenetic analyses were performed using only the SNVs and variants from PE/PPE/PGRS were removed using bedtools intersect using mask regions provided in Snippy (39). Merged multi-sample vcf files were converted to a fasta file containing the genotypes using vcf2fasta tool. Construction of a phylogenetic tree was performed using IQ-TREE2 (24). We have restricted model selection to GTR models and automatic model selection was done with ModelFinder Plus implementation. SH-aLRT test and 1000 bootstrap replicates were computed using UFBoot2 implementation in the tool (40, 41).

### PCA of Indian and global tuberculosis isolates

For PCA of tuberculosis samples we downloaded a collection of more than 2500 GMTV database and merged all the files with all the Indian tuberculosis genomic variant datasets and an aligned SNV fasta file was generated. Then for pairwise distance among the samples were calculated using snp-dist tool. Principal components were calculated and plotted using PCAtools R package.

### Statistical analysis

Except for the mentioned cases, the rest of the statistical tests were performed using base R functions and plotted using the GGPUBR wrapper of the ggplot2 package.

## Data & code availability

All the whole genome sequencing raw datasets will be uploaded to NCBI SRA repository upon acceptance of this manuscript for publication. The code used for analysis and statistical tests will be made available after publication of this manuscript.

## Author contributions

AG, SKR planned the study and co-ordinated the data analysis and sample collection with ICMR-RMRC team. AG, VKB performed the DNA extraction and library preparation of Mtb samples. HB, DD provided access to BSL-3 facility, Mtb cultures and respective metadata. AG wrote the manuscript SKR, DD and SP edited the manuscript.

## Acknowledgments

AG is supported by ICMR-SRF fellowship, VKB is supported by GenomeIndia fellowship. To perform the study SKR received the funding from DBT-ILS intramural grant.

---

**Supplementary Figure 1** A) Age distribution of patients enrolled for the study. B) Fractional distribution of MTBC detected in sequenced samples. C) INDEL length and count distribution, where negative lengths represents deletions and positive lengths insertions. D) Distribution of deletions detected in using Delly2 pipeline. E) Functional annotation of single nucleotide variants.

**Supplementary Figure 2** A) Hierarchical clustering of pair-wise SNP distances annotated with lineage state and drug resistance phenotypes.

## References

1. World Health Organization. 2021. Global Tuberculosis Report 2021Global Tuberculosis Report.

2. World Health Organization (WHO). 2020. WHO report on TB 2020Who.

3. Ndjeka N, Schnippel K, Master I, Meintjes G, Maartens G, Romero R, Padanilam X, Enwerem M, Chotoo S, Singh N, Hughes J, Variava E, Ferreira H, Te Riele J, Ismail N, Mohr E, Bantubani N, Conradie F. 2018. High treatment success rate for multidrug-resistant and extensively drug-resistant tuberculosis using a bedaquiline-containing treatment regimen. Eur Respir J 52.

4. Ryu YJ. 2015. Diagnosis of pulmonary tuberculosis: recent advances and diagnostic algorithms. Tuberc Respir Dis (Seoul) 78:64–71.

5. Manson AL, Abeel T, Galagan JE, Sundaramurthi JC, Salazar A, Gehrmann T, Shanmugam SK, Palaniyandi K, Narayanan S, Swaminathan S, Earl AM. 2017. Mycobacterium tuberculosis whole genome sequences from Southern India suggest novel resistance mechanisms and the need for region-specific diagnostics. Clin Infect Dis 64:1494–1501.

6. Manson AL, Abeel T, Galagan JE, Sundaramurthi JC, Salazar A, Gehrmann T, Shanmugam SK, Palaniyandi K, Narayanan S, Swaminathan S, Earl AM. 2017. Mycobacterium tuberculosis Whole Genome Sequences From Southern India Suggest Novel Resistance Mechanisms and the Need for Region-Specific Diagnostics. Clin Infect Dis 64:1494–1501.

7. Advani J, Verma R, Chatterjee O, Pachouri PK, Upadhyay P, Singh R, Yadav J, Naaz F, Ravikumar R, Buggi S, Suar M, Gupta UD, Pandey A, Chauhan DS, Tripathy SP, Gowda H, Prasad TSK. 2019. Whole genome sequencing of Mycobacterium tuberculosis clinical isolates from India reveals genetic heterogeneity and region-specific variations that might affect drug susceptibility. Front Microbiol 10:1–15.

8. Chatterjee A, Nilgiriwala K, Saranath D, Rodrigues C, Mistry N. 2017. Whole genome sequencing of clinical strains of Mycobacterium tuberculosis from Mumbai, India: A potential tool for determining drug-resistance and strain lineage. Tuberculosis 107:63–72.

9. Singh J, Sankar MM, Kumar P, Couvin D, Rastogi N, Singh S, Katoch VM, Chauhan DS, Katoch K, Rodrigues C, Lakshmi V, Taori GM, Daginawala HF, Singh R, Bhattacharya B, Choudhury B, Singh N, Devi U, Swaminathan S. 2015. Genetic diversity and drug susceptibility profile of Mycobacterium tuberculosis isolated from different regions of India. J Infect 71:207–219.

10. Coll F, Phelan J, Hill-Cawthorne GA, Nair MB, Mallard K, Ali S, Abdallah AM, Alghamdi S, Alsomali M, Ahmed AO, Portelli S, Oppong Y, Alves A, Bessa TB, Campino S, Caws M, Chatterjee A, Crampin AC, Dheda K, Furnham N, Glynn JR, Grandjean L, Minh Ha D, Hasan R, Hasan Z, Hibberd ML, Joloba M, Jones-López EC, Matsumoto T, Miranda A, Moore DJ, Mocillo N, Panaiotov S, Parkhill J, Penha C, Perdigão J, Portugal I, Rchiad Z, Robledo J, Sheen P, Shesha NT, Sirgel FA, Sola C, Oliveira Sousa E, Streicher EM, Helden P Van, Viveiros M, Warren RM, McNerney R, Pain A, Clark TG. 2018. Genome-wide analysis of multi-and extensively drug-resistant Mycobacterium tuberculosis. Nat Genet 50:307–316.

11. Comas I, Coscolla M, Luo T, Borrell S, Holt KE, Kato-Maeda M, Parkhill J, Malla B, Berg S, Thwaites G, Yeboah-Manu D, Bothamley G, Mei J, Wei L, Bentley S, Harris Niemann S, Diel R, Aseffa A, Gao Q, Young D, Gagneux S. 2013. Out-of-Africa migration and Neolithic coexpansion of Mycobacterium tuberculosis with modern humans. Nat Genet 45:1176–1182.

12. Chen ML, Doddi A, Royer J, Freschi L, Schito M, Ezewudo M, Kohane IS, Beam A, Farhat M. 2019. Beyond multidrug resistance: Leveraging rare variants with machine and statistical learning models in Mycobacterium tuberculosis resistance prediction. EBioMedicine 43:356–369.

13. Yang Y, Niehaus KE, Walker TM, Iqbal Z, Walker AS, Wilson DJ, Peto TEA, Crook DW, Smith EG, Zhu T, Clifton DA. 2018. Machine learning for classifying tuberculosis drug-resistance from DNA sequencing data. Bioinformatics 34:1666–1671.

14. Yang Y, Walker TM, Walker AS, Wilson DJ, Peto TEA, Crook DW, Shamout F, CRyPTIC Consortium, Zhu T, Clifton DA. 2019. DeepAMR for predicting co-occurrent resistance of Mycobacterium tuberculosis. Bioinformatics 35:3240–3249.

15. Chatterjee A, Nilgiriwala K, Saranath D, Rodrigues C, Mistry N. 2017. Whole genome sequencing of clinical strains of Mycobacterium tuberculosis from Mumbai, India: A potential tool for determining drug-resistance and strain lineage. Tuberculosis 107:63–72.

16. Gutierrez MC, Ahmed N, Willery E, Narayanan S, Hasnain SE, Chauhan DS, Katoch VM, Vincent V, Locht C, Supply P. 2006. Predominance of ancestral lineages of Mycobacterium tuberculosis in India. Emerg Infect Dis 12:1367–74.

17. Merker M, Barbier M, Cox H, Rasigade JP, Feuerriegel S, Kohl TA, Diel R, Borrell S, Gagneux S, Nikolayevskyy V, Andres S, Nübel U, Supply P, Wirth T, Niemann S. 2018. Compensatory evolution drives multidrug-resistant tuberculosis in central Asia. Elife 7:1–31.

18. Manson AL, Cohen KA, Abeel T, Desjardins CA, Armstrong DT, Barry CE, Brand J, Chapman SB, Cho SN, Gabrielian A, Gomez J, Jodals AM, Joloba M, Jureen P, Lee JS, Malinga L, Maiga M, Nordenberg D, Noroc E, Romancenco E, Salazar A, Ssengooba W, Velayati AA, Winglee K, Zalutskaya A, Via LE, Cassell GH, Dorman SE, Ellner J, Farnia P, Galagan JE, Rosenthal A, Crudu V, Homorodean D, Hsueh PR, Narayanan S, Pym AS, Skrahina A, Swaminathan S, Van Der Walt M, Alland D, Bishai WR, Cohen T, Hoffner S, Birren BW, Earl AM. 2017. Genomic analysis of globally diverse Mycobacterium tuberculosis strains provides insights into the emergence and spread of multidrug resistance. Nat Genet 49:395–402.

19. Wattam AR, Abraham D, Dalay O, Disz TL, Driscoll T, Gabbard JL, Gillespie JJ, Gough R, Hix D, Kenyon R, Machi D, Mao C, Nordberg EK, Olson R, Overbeek R, Pusch GD, Shukla M, Schulman J, Stevens RL, Sullivan DE, Vonstein V, Warren A, Will R, Wilson MJC, Yoo HS, Zhang C, Zhang Y, Sobral BW. 2014. PATRIC, the bacterial bioinformatics database and analysis resource. Nucleic Acids Res 42:D581–91.

20. Anyansi C, Keo A, Walker BJ, Straub TJ, Manson AL, Earl AM, Abeel T. 2020. QuantTB-A method to classify mixed Mycobacterium tuberculosis infections within whole genome sequencing data. BMC Genomics 21.

21. Coll F, McNerney R, Guerra-Assunção JA, Glynn JR, Perdigão J, Viveiros M, Portugal I, Pain A, Martin N, Clark TG. 2014. A robust SNP barcode for typing Mycobacterium tuberculosis complex strains. Nat Commun 5:4–8.

22. Freschi L, Vargas R, Husain A, Kamal SMM, Skrahina A, Tahseen S, Ismail N, Barbova A, Niemann S, Cirillo DM, Dean AS, Zignol M, Farhat MR. 2021. Population structure, biogeography and transmissibility of Mycobacterium tuberculosis. Nat Commun 12.

23. Devi KR, Pradhan J, Bhutia R, Dadul P, Sarkar A, Gohain N, Narain K. 2021. Molecular diversity of Mycobacterium tuberculosis complex in Sikkim, India and prediction of dominant spoligotypes using artificial intelligence. Sci Rep 11:7365.

24. Minh BQ, Schmidt HA, Chernomor O, Schrempf D, Woodhams MD, Von Haeseler A, Lanfear R, Teeling E. 2020. IQ-TREE 2: New Models and Efficient Methods for Phylogenetic Inference in the Genomic Era. Mol Biol Evol 37:1530–1534.

25. Chernyaeva EN, Shulgina M V., Rotkevich MS, Dobrynin P V., Simonov SA, Shitikov EA, Ischenko DS, Karpova IY, Kostryukova ES, Ilina EN, Govorun VM, Zhuravlev VY, Manicheva OA, Yablonsky PK, Isaeva YD, Nosova EY, Mokrousov I V., Vyazovaya AA, Narvskaya O V., Lapidus AL, O’Brien SJ. 2014. Genome-wide Mycobacterium tuberculosis variation (GMTV) database: A new tool for integrating sequence variations and epidemiology. BMC Genomics 15.

26. Bradley P, Gordon NC, Walker TM, Dunn L, Heys S, Huang B, Earle S, Pankhurst LJ, Anson L, De Cesare M, Piazza P, Votintseva AA, Golubchik T, Wilson DJ, Wyllie DH, Diel R, Niemann S, Feuerriegel S, Kohl TA, Ismail N, Omar S V., Smith EG, Buck D, McVean G, Walker AS, Peto TEA, Crook DW, Iqbal Z. 2015. Rapid antibiotic-resistance predictions from genome sequence data for Staphylococcus aureus and Mycobacterium tuberculosis. Nat Commun 6.

27. Farhat MR, Jesse Shapiro B, Kieser KJ, Sultana R, Jacobson KR, Victor TC, Warren RM, Streicher EM, Calver A, Sloutsky A, Kaur D, Posey JE, Plikaytis B, Oggioni MR, Gardy JL, Johnston JC, Rodrigues M, C Tang PK, Kato-Maeda M, Borowsky ML, Muddukrishna B, Kreiswirth BN, Kurepina N, Galagan J, Gagneux S, Birren B, Rubin EJ, Lander ES, Sabeti PC, Murray M. 2013. Genomic analysis identifies targets of convergent positive selection in drug-resistant Mycobacterium tuberculosis. Nat Publ Gr 45.

28. Vargas R, Freschi L, Marin M, Epperson LE, Smith M, Oussenko I, Durbin D, Strong M, Salfinger M, Farhat MR. 2021. In-host population dynamics of mycobacterium tuberculosis complex during active disease. Elife 10:1–37.

29. Ektefaie Y, Dixit A, Freschi L, Farhat MR. 2021. Globally diverse Mycobacterium tuberculosis resistance acquisition: a retrospective geographical and temporal analysis of whole genome sequences. The Lancet Microbe 2:e96–e104.

30. Hussain T, Tripathy SS, Das S, Satapathy P, Das D, Thomas B, Pati S. 2020. Prevalence, risk factors and health seeking behaviour of pulmonary tuberculosis in four tribal dominated districts of Odisha: Comparison with studies in other regions of India. PLoS One 15.

31. Rao VG, Bhat J, Yadav R, Sharma RK, Muniyandi M. 2019. Declining tuberculosis prevalence in Saharia, a particularly vulnerable tribal community in Central India: evidences for action. BMC Infect Dis 19:180.

32. Vargas R, Freschi L, Marin M, Epperson LE, Smith M, Oussenko I, Durbin D, Strong M, Salfinger M, Farhat MR. 2021. In-host population dynamics of Mycobacterium tuberculosis complex during active disease. Elife 10.

33. Li H. 2013. Aligning sequence reads, clone sequences and assembly contigs with BWA-MEM.

34. Walker BJ, Abeel T, Shea T, Priest M, Abouelliel A, Sakthikumar S, Cuomo CA, Zeng Q, Wortman J, Young SK, Earl AM. 2014. Pilon: An integrated tool for comprehensive microbial variant detection and genome assembly improvement. PLoS One 9.

35. Danecek P, Bonfield JK, Liddle J, Marshall J, Ohan V, Pollard MO, Whitwham A, Keane T, McCarthy SA, Davies RM, Li H. 2021. Twelve years of SAMtools and BCFtools. Gigascience 10.

36. Cingolani P, Platts A, Wang LL, Coon M, Nguyen T, Wang L, Land SJ, Lu X, Ruden DM. 2012. A program for annotating and predicting the effects of single nucleotide polymorphisms, SnpEff: SNPs in the genome of Drosophila melanogaster strain w1118; iso-2; iso-3. Fly (Austin) 6:80–92.

37. Cingolani P, Patel VM, Coon M, Nguyen T, Land SJ, Ruden DM, Lu X. 2012. Using Drosophila melanogaster as a model for genotoxic chemical mutational studies with a new program, SnpSift. Front Genet 3.

38. Xia E, Teo Y-Y, Ong RT-H. 2016. SpoTyping: fast and accurate in silico Mycobacterium spoligotyping from sequence reads. Genome Med 8:19.

39. Quinlan AR, Hall IM. 2010. BEDTools: A flexible suite of utilities for comparing genomic features. Bioinformatics 26:841–842.

40. Kalyaanamoorthy S, Minh BQ, Wong TKF, von Haeseler A, Jermiin LS. 2017. ModelFinder: fast model selection for accurate phylogenetic estimates. Nat Methods 14:587–589.

41. Hoang DT, Chernomor O, von Haeseler A, Minh BQ, Vinh LS. 2018. UFBoot2: Improving the Ultrafast Bootstrap Approximation. Mol Biol Evol 35:518–522.

