## Supplementary figures and images for "Genotypic and phenotypic diversity of the multidrug-resistant *Mycobacterium tuberculosis* strains from eastern India"

### Supplementary Figure 1

#### Supplementary Figure 1

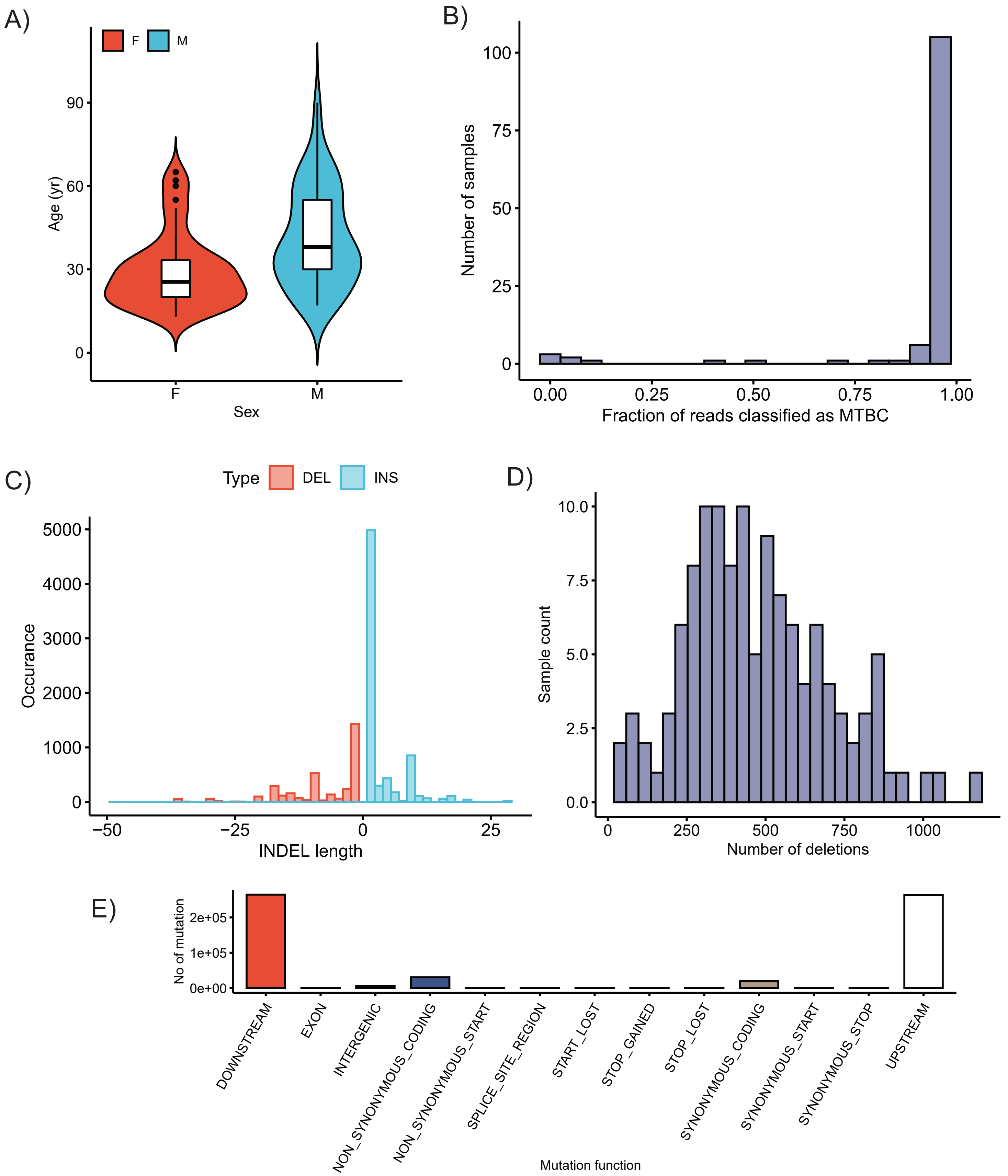

### Supplementary Figure 2

#### Supplementary Figure 2

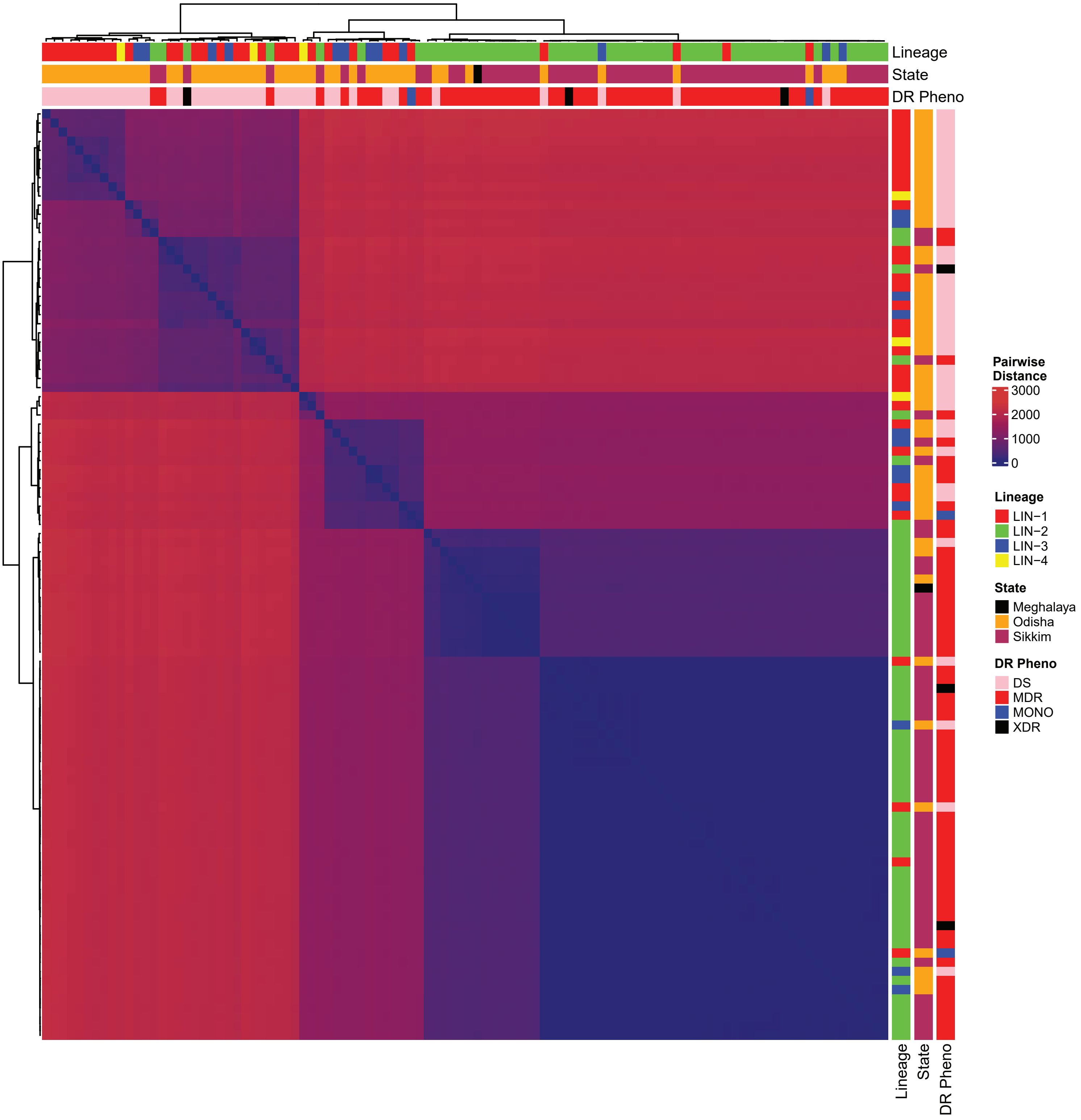
